# Integrated-omics profiling unveils the disparities of host defense to ECM scaffolds during wound healing in aged individuals

**DOI:** 10.1101/2023.12.26.573379

**Authors:** Shuai-dong Chen, Chen-yu Chu, Chen-bing Wang, Yi Man

**Affiliations:** State Key Laboratory of Oral Diseases & National Center for Stomatology & National Clinical Research Center for Oral Diseases & Department of Oral Implantology, West China Hospital of Stomatology, Sichuan University, Chengdu 610041, Sichuan, China

## Abstract

Extracellular matrix (ECM) scaffold membranes have exhibited promising potential to better the outcomes of wound healing by creating a regenerative microenvironment around. However, when compared to the application in younger individuals, the performance of the same scaffold membrane in promoting re-epithelialization and collagen deposition was observed dissatisfying in aged mice. To comprehensively elucidate the mechanisms underlying this age-related disparity, we conducted an integrated analysis, combing single-cell RNA sequencing (scRNA-Seq) with spatial transcriptomics, to explore the complex cellular niches surrounding the ECM scaffolds. Through intergroup comparative analysis and cell-cell communication, we identified and characterized the senescent SPP1+ macrophages may impede the activation of the type L immune response, thus inhibiting the repair ability of epidermal cells and fibroblasts around the ECM scaffolds. These findings contribute to a deeper understanding of biomaterial applications in varied physiological contexts, thereby paving the way for the development of precision-based biomaterials tailored specifically for aged individuals in future therapeutic strategies.

## Background

Wound healing is a complex process that receives great attention. Following injury, the wound undergoes a series of dynamic and overlapping phases to restore the tissue defects, including the early hemostasis and acute inflammatory responses, followed by the re-epithelialization of the epidermis and the fibrous repair of the dermis in later stages^1^. Based on investigations on the mechanisms underlying wound healing process, the faster wound closure and superior healing outcomes are expected. Therefore, biomaterials based on extracellular matrix (ECM), assembled from collagen and proteoglycans, have been developed and demonstrated promising effects in promoting wound regeneration^2,3^.

Despite the various components and forms, ECM-based biomaterials primarily derive from exogenous proteins^4^. Upon implantation, these biomaterials, recognized as foreign bodies, often recruit neutrophils and monocytes, facilitating the clearance and phagocytosis of damaged cells and invaded bacteria in the early stages of local implant sites and reducing the risk of wound infection^5^. In the later stages, the recruited monocytes can differentiate into macrophages with different polarization status, secreting growth factors that promote the wound repair process. Additionally, ECM-based biomaterials with the immunogenicity can be recognized and presented to the adaptive immune cells, genera^6,7^ting the type II immune response to facilitate the wound repair and the neogenesis of skin appendages^8–10^. Thus, the ECM-based biomaterial may serve as a promising approach to better the wound healing.

Nowadays, ECM-based biomaterials have been evaluated the regenerative ability mostly in young individuals. However, the alterations in skin structure and cellular functionality of the aged skin, pose the significant difference of wound healing process in aged individuals. Current evidence supports that the chemotaxis and function of neutrophils as well as macrophages are greatly diminished in aged individuals, related to the increased local bacterial colonization, and delayed wound closure^11^. In the epithelia repair phase, epidermal stem cells with reduced hemidesmosome components and glycosaminoglycans^6,7^, exhibit increased expression of differentiation markers such as Krt10 and downregulated proliferative capacity^12^, impairing wound re-epithelialization in aged wounds. Besides, the aged fibroblasts upregulate the differentiation potential into adipocytes while downregulated collagen-producing capacity, resulting in unsatisfying repair outcomes^13,14^. In addition to the wound healing difference, aging also leads to substantial alterations in immune cells, particularly macrophages^15,16^, not only causing significant changes in the composition and polarization status around implants but also influencing the recognition and functional responses of immune system to materials.

Hence, these age-related alterations may exhibit risks in the application of ECM-based materials for wound repair in the elderly. Within the guidance of precision medicine, the performance of the same ECM scaffold and the interaction with the aged individuals remains to be clarified^17–19^. However, the host response to the biomaterials is exceedingly complex, influenced not only by various cell types, signaling pathways, and environmental factors but also by the spatial positioning of cell compositions, posing a challenge in accurately deciphering the crosstalk between biomaterials and the host^20,21^. To solve that, we integrate the single-cell RNA sequencing (scRNAseq) with spatial transcriptomics (ST) to dissect the surrounding cells at a high-resolution singlecell level as well as to establish their functional niches around, thus gaining a better understanding of the disparities of microenvironment around the ECM scaffold membranes, which would provide new insights into the custom design and evaluation of the wound healing biomaterials in future therapeutic strategies.

## Results

### 1. ECM scaffold did not enhance wound healing process in aged mice

To begin with, mice were categorized into young and aged groups and conducted with fullthickness wounds (6mm in diameter). Then, we placed ECM scaffold membranes in the wounds in ECM group while control group received with normal saline in each age group as in Fig. 1b.

**Figure1.**
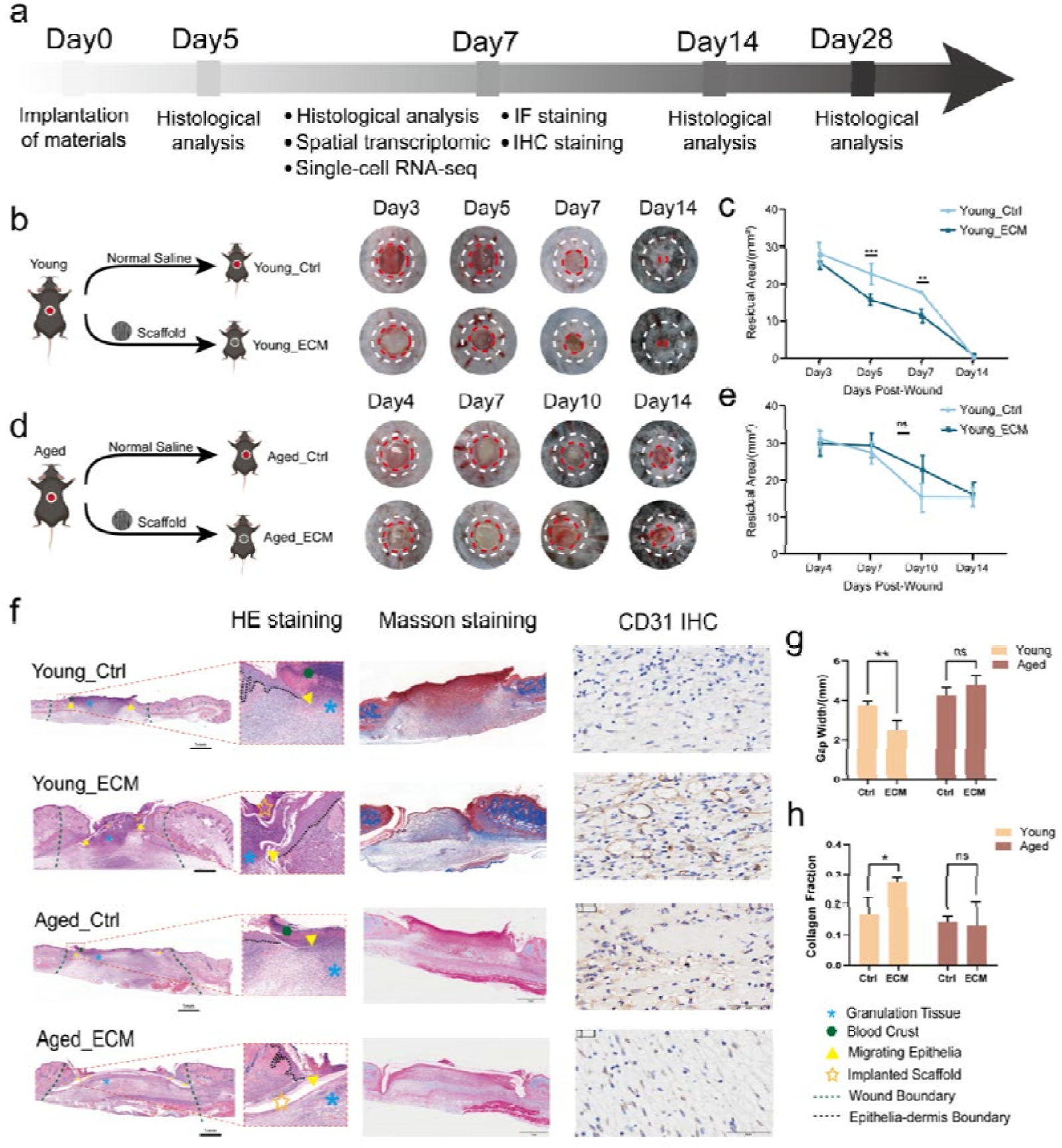
The different effect of ECM scaffolds on wound healing in young and aged mice. (a) Workflow for evaluating the wound healing outcomes in young and aged mice. (b-c) Schematic diagram for the surgical process for full-thickness wound model and the residual wound area at 3, 5, 7, 14 days in young mice. (d-e) Schematic diagram for the surgical process for full-thickness wound model and the residual wound area at 4, 7, 10, 14 days in aged mice. (f) The histological results for H&E as well as Masson staining and the IHC results for CD31 of wounds at POD7 in Control group and ECM group of young and aged mice. (g) Semiquantitative evaluation of gap width to H&E staining results in Control group and ECM group of young and aged mice (n=3 for each group). (h) Semiquantitative evaluation of collagen deposition to Masson staining results in Control group and ECM group of young and aged mice (n=3 for each group). **P<0.01, *P<0.05 and ns/no significance by Student’s t-test for data in (g) and (h), and TWO-WAY ANOVA for data in (c) and (e).

The workflow to evaluate the potential of ECM scaffolds membrane in wound healing in young and aged mice was summarized in Fig.1a. In young mice, control group (Young_Ctrl) exhibited a steady wound closure with over half residual aera on postoperative day (POD) 7 while wound closure in the ECM group (Young_ECM) was accelerated and superior to Young_Control on POD 5 and POD 7 (Fig. 1c), in accordance with the histological results of epithelial gap width to access the re-epithelization (Extended Data. Fig1a, b). Although both groups achieved visible and complete wound closure in young wounds on POD 14, the hair follicles neogenesis was observed exclusively in Young_ECM group (Extended Data. Fig1c), demonstrating the ability of ECM scaffolds to promote epidermis functional regeneration in the whole wound healing process. Conversely, in aged mice, the control group (Aged_Control) did not display a distinct tendency for wound closure until POD 7 and nearly half of the wound remained incompletely closed on POD 14 (Fig. 1d) aligning with the previous reports^22^. After the implantation of scaffold, ECM group showed no significant acceleration of epithelial repair in any timepoint both in would closure and epithelium gap width, even indicating a tendency to impair wound healing on POD7 (Fig. 1e). Thus, the ECM scaffold membrane to promote functional epidermis regeneration in young mice did not show the same effect in aged wounds (Fig. 1g).

Subsequent, Masson’s trichrome staining was employed to evaluate local collagen fiber deposition and induction of dermal repair of the ECM scaffold. On POD 7, below the crust of the necrotic cell debris, the collagen fibers were irregularly arranged in the granulation tissue in Young_Control group while Young_ECM group showed the increased collagen fiber deposition in a more stratified and organized pattern (Fig. 1f, h). On POD 14, due to the activation of fibroblast and wound remodeling, deposited collagen was future mature in Young_Control and more abutment and similar to the normal dermis structure in Young_ECM (Extended Data. Fig 1.e, f). However, the collagen fibers in the granulation tissue were not significantly different between Aged_Control and Aged_ECM groups on POD 7 or PO7 14. Besides the collagen deposition, angiogenesis, crucial for the maintenance of newly formed granulation tissue^23^, was evaluated through the immunohistochemistry (IHC) staining for CD31 to further assess dermal repair. Young_ECM group shows the more abundant and widespread vascular endothelial staining than Control group while the Aged_ECM group still shows the inability to angiogenesis both on POD 7 and POD 14 (Fig. 1f, Extended Data. Fig 1.f-h). Therefore, in the aged mice, the ECM scaffold was unable to promote the formation of granulation tissue and collagen repair in the dermis.

In summary, the ECM scaffold membrane effectively facilitated the epidermal functional regeneration and the dermal repair process, with the faster wound closure and more abundant and regular collagen deposition and angiogenesis while failed to facilitate comprehensive wound repair processes in the aged wounds.

### 2. Acute inflammation persists around the ECM scaffold in the aged wounds

Our previous reports demonstrate the ability to enhance wound healing of implanted ECM scaffold membrane can be attributed to local regenerative immune microenvironment^24^. Then, we speculate the reason of the adverse performance of ECM scaffold in aged wounds may be the alteration of the microenvironment. Since the POD7 is a crucial timepoint to transit from the innate to adaptive immunity, we combined scRNA-seq with ST to dissect the different host defense to the scaffolds in Young_ECM and Aged_ECM as in Fig. 2a.

**Figure2.**
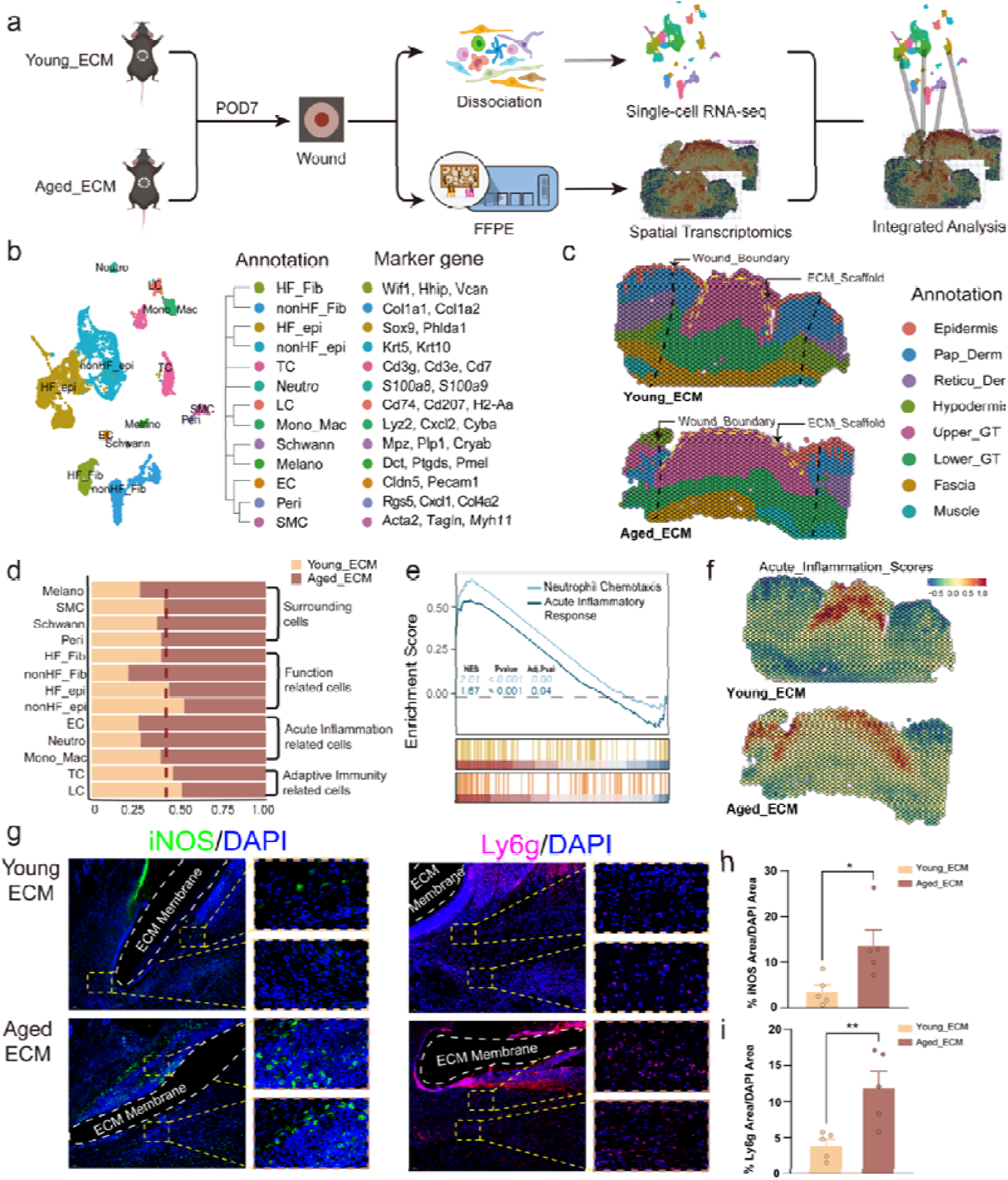
The acute inflammation rather than type □ immune response around ECM scaffold membrane in aged wounds. (a) Schematic diagram for generating the scRNA-seq and spatial transcriptomics data from ECM mediated wound healing at POD7 in young and aged mice. (b) The UMAP reduction results reveals 13 different groups in wound. The marker genes are listed on the right of the annotation. (c) The annotated clustering results for spatial transcriptomic based on the anatomic structure of the wounds. (d) The proportion of each cell type of the whole wounds and their related wound healing process. (e) The upregulated results for Gene Sets Enrichment Analysis (GSEA) in ECM scaffolds treated aged wounds. (f) The AddModule scores of acute inflammation term in spatial transcriptomics expression. (g) The IF results for iNOS and Ly6g in Young_ECM and Aged_ECM at POD7. (h-i) The semiquantitative analysis of iNOS and Ly6g fluorescence area in (g) respectively (n=3 for each group). **P<0.01 and *P<0.05 by Student’s t-test for data in (h) and (i).

After quality control, 52 distinct clusters encompassing 13 cell types were identified via UMAP reduction of Seurat (Fig. 2b). We annotated these cell types by known genes: HF-unrelated epidermal cells /nonHF_Epi (*Ly6a+, Krt5+, Krt10+*)^25^, HF-related epidermal cells /HF_Epi (*Ly6a-, Sox9+, Lgr5+*)^26^, HF-related fibroblasts /HF_Fib (*Corin+, Wif1+, Fgf7+*), HF-unrelated fibroblasts /nonHF_Fib (*Igfbp3+, Col1a1+, Col15a1+*), Langerhans cells /LC (*Cd207+, Cd74+*), Monocytederived macrophages /Mono_DC (*Lyz2+, Cd74+*), Neutrophils /Neutro (*S100a8+, Retnlg+*), T cells /TC (*Cd3d+, Trac+, Trdc+*), Melanocytes /Melano (*Dct+, Ptgds+*), Schwann cells / Schwann (*Mpz+, Plp1+*), Endothelial cells /EC (*Pecam1+, Cldn5+*), Pericytes /Peri (*Rgs5+, Sparcl1+*), and Smooth muscle cells /SMC (*Tagln+, Acta2+*). The spatial transcriptomics results were clustered using unsupervised clustering and annotated based on the anatomic structure as Epidermis, Papillary dermis, Reticular dermis, Dermal sublayers, Granulation tissue, Fascia, and Muscle (Fig. 2c).

We then discovered higher proportions of acute inflammation associated neutrophils and macrophages in the Aged_ECM, and the more abundant adaptive immune cells such as Langerhans cells and T cells in Young_ECM (Fig. 2d). In accordance, Young_ECM upregulated genes associated with dendritic cells and T cells (H2-Ab1, H2-Aa, Cd3g, Cxcl14) (Extended Data. Fig 2.d), whereas Aged_ECM upregulated genes associated with neutrophils and macrophages (S100a8, S100a9, Tnfaip6) and corresponding GO enrichment of “acute inflammatory response” and “neutrophil chemotaxis”(Fig 2.e). Later, we projected the enriched acute inflammation gene sets to the spatial transcriptomic slices through AddmoduleScore function in Seurat, which reveals not only acute inflammation beneath the scaffold but also a more extensive range around the lower dermis in Aged_ECM (Fig 2.f). To validate this, we used immunofluorescence (IF) staining for iNOS^27^ and ly6g^28^ to mark macrophages and neutrophils respectively and found a higher proportion of innate response cell expression in the granulation tissue below the ECM scaffold and papillary dermis around Aged_ECM (Fig 2.g-i).

During the acute inflammation stage after ECM scaffold implantation, recruited neutrophils and macrophages are expected to play a defense role in the clearance of necrotic debris and invaded pathogens. Hence, we analyzed the intergroup functional differences of neutrophils and macrophages and found these innate immune cells in the Aged_ECM group upregulated inflammatory cells chemotaxis and intrinsic apoptosis while downregulated the defense against foreign pathogens, phagocytosis of apoptotic cells, and regulation of the adaptive immune responses (Extended Data. Fig 3. a-d). Moreover, the Aged_ECM group upregulated the expression of proinflammatory cytokines like IL-1β^29,30^ and TNF-^31^, responding to tissue damage and creating a local positive feedback amplification of acute inflammation, recruiting more inflammatory cells to the area(Extended Data. Fig 3. e-g). Therefore, the defects in the proper function in innate immune cells around Aged_ECM group led to a widespread and sustained acute inflammatory response, affecting the transition towards a regenerative microenvironment in the aged wounds.

**Figure3.**
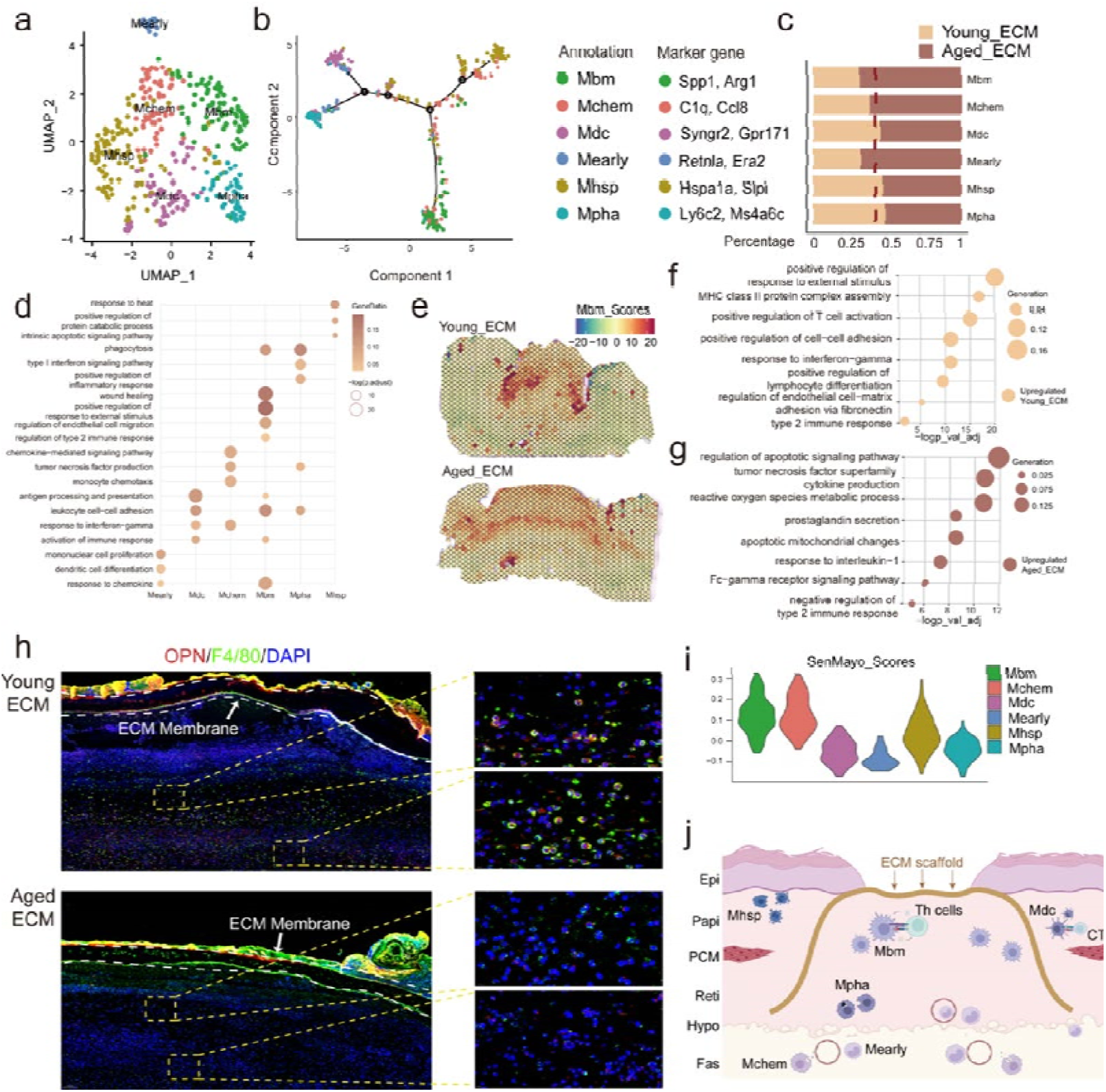
The heterogeneity of macrophages and the intergroup difference of Mbm. (a-b) The UMAP and the DDTree reduction results for the subclustering of Mono_Mac in Fig2b. The annotations and marker genes are listed in the right. (c-d) The proportion and Gene Ontology (GO) enrichment results of each cell subtype of the macrophages in Young_ECM and Aged_ECM. (e) The AddModule score projection of Mbm-specific genes in scRNA-seq results. (f-g) The upregulated GO enrichment terms of Mbm subtypes in Young_ECM and Aged_ECM, respectively. (h) The Immunofluorescence double staining for OPN/Spp1 and F4/80 results shows the reduced the double positive cells in Aged_ECM. (i) The SenMayo gene set scores for subtypes of macrophages. (j) The schematic diagram for the spatial location of each subtype for macrophage in the ECM mediated wound healing process.

### 3. The heterogeneity of macrophages responds differently to the ECM scaffold

Besides the preliminary discovery of acute inflammation around scaffolds in Aged_ECM and the intergroup difference in related cell types, cells of the same type can exhibit different polarization states and functional responses toward the ECM scaffolds, especially in macrophages. Thus, we subset macrophages, a bridge connecting the acute inflammation with the adaptive immune response and run the sub-cluster analysis of their functional heterogeneity. Unlike the traditional M1/M2 classification system, macrophages around the ECM scaffold can be classified functionally into six subclusters: Mpha (*Ms4a+, Ly6c2+*), Mearly (*Retnla+, Ear2+*)^32^, Mchem (*Ccl8+, Ccl7+*), Mhsp (*Hspa1a+, Hspa1b+*)^33,34^, Mdc (*H2-Aa+, Cd74+*), Mbm (*Spp1+, Arg1+*)^35^ (Fig 3.a, Extended Data. Fig 4. a).

Among these types, Mhsp, expressing heat shock protein associated genes, is distributed in the epidermis and papillary dermis around the wound. Since the upregulated genes are not significant and only involve functions such as protein degradation and apoptosis, Mhsp may act as the decreasing pro-inflammatory macrophages during the resolution of acute inflammation (Fig 3.c, Extended Data. Fig 4. k). Conversely, Mpha in the lower the granulation tissue, is responsible for the phagocytosis of apoptotic cells and exaggeration of local inflammation by secreting proinflammatory factor TNF-α (Extended Data. Fig 4. j). Additionally, Mchem, located in the fascia below the granulation tissue and reticular dermis around, can respond to the increased IFN-γ and recruit more monocytes to the ECM scaffold through chemotactic factors such as Ccl8^36,37^ and Ccl7^38^, facilitating the later wound repair process(Extended Data. Fig 4. g). *Retnla+Ear2+* Mearly, scattered predominantly around the lower granulation tissue and reticular dermis, co-localizes with endothelia cells in blood vessels (Extended Data. Fig 4. f, i). It not only responds to chemotactic factors but also upregulates functions related to monocyte proliferation, indicating potential the role of precursor cell status from circulation, as previously described^32^. Furthermore, Mearly demonstrates the potential for differentiation into dendritic cells and indeed as the same brunch in DDTree reduction (Fig 3.b), within its spatial vicinity, antigen-presenting Mdc expressing MHC-II molecules are discovered, which may stimulate the activation of adaptive immune responses in later stages. Finally, *Spp1+, Arg1+* Mbm, previously identified as the responder to biomaterials^35^, was abundant beneath the scaffold and exhibited GO enrichment associated with reactions to external stimulus (Fig 3.e). Besides, Mbm exhibited multiple biological functions, not only enhancing phagocytic abilities to clear cell debris but also regulating type II immune responses and contributing to wound healing (Fig 3. d, Extended Data. Fig 4. b, e). The spatial anchor and the function analysis together suggest the crucial role of Mbm in the interaction between the ECM scaffold and the host implanted (Fig 3.h).

Subsequently, we explored intergroup differences among macrophage subtypes to explain the distinct performance of ECM scaffolds in aged wounds. The differential expressed genes and functional enrichment of Mhsp, Mpha, and Mchem did not show significant intergroup differences, only exhibiting disparities in spatial localization and abundance, in accordance with the sustained acute inflammation around scaffolds in Aged_ECM. On the contrary, Mearly remained spatially stable around the ECM scaffold, but gene enrichment results indicated downregulation of proliferation ability, implying impaired differentiation towards macrophages entering the wound in Aged_ECM (Extended Data. Fig 4. c).

However, Mdc and Mbm exhibited not only the reduced abundance and distribution around the scaffold but also the downregulation of intrinsic functions involved in the interaction with adaptive immune response. Mdc shows decreased expression in the reticular dermis around the ECM scaffold and weakening abilities to regulate cytotoxicity in aged wounds, suggesting its association with functional obstacles in CTLs and the clearance of damaged cells (Extended Data. Fig 4. d). While Mbm, closer to the scaffold spatially, also showed reduced abundance around scaffolds but its diminished abilities to regulate the adaptive immune response were related to the helper T cells, which helps explain why the type II immune responses were not provoked by ECM scaffolds in aged individuals as in young wounds (Fig 3. f-g). When aging, cellular senescence often predominates the reason for the declining systemic function^39,40^. To figure out whether these intergroup differences of Mdc and Mbm are caused by cellular senescence, we evaluated the senescence scores of different subtypes using the SenMayo gene set^41^ and found that Mbm exhibited the highest senescence scores (Fig 3. i), suggesting the intergroup differences in the function of Mbm functionality might be more related to intrinsic cellular senescence^42^.

Therefore, we believe that following the implantation of the ECM scaffold in aged group wounds, different macrophage subtypes will exert unique functions in distinct spatial locations (Fig 3j), and the differences in their abundance are related to the persisted acute inflammation. Among them, Mbm, significantly affected by cellular senescence, may play a central role in the altered microenvironment around the ECM scaffold in Aged_ECM.

### 3. Age-related alterations in regenerative Th cells are less affected by cellular senescence

After exploring various functional and intergroup difference into the innate immune response related macrophages, our focus shifted to T cells given the interconnection between adaptive immune responses around the scaffold and functional wound healing^43,44^. Initially, we segregated T cells in the first reduction level into αβ T cells and γδ T cells based on *Trdc* expression. As adaptive immune responses are primarily mediated by αβ T cells, we further subset the αβ T cells for secondary dimensional reduction clustering and identified a total of 7 functional subclusters: Naïve helper T cells/Th_naïve (*Cd4+, Cd28+*), Memory T cells /Tm (*Hlf+, Ccr7+*)^45^, Cytotoxic T cells /CTL (*Cd8a+, Cd8b1+*), IL-17 producing T cells /T17 (*Il17a+, Il17f+*), Nature killer cells/NKT (*Nkg2d+, Gzmc+*), Regulatory T cells /Treg (Foxp3+, Ctla4+), Innate Lymphoid CelIs /ILC (*Sox9+, Gata3+*)^46^ (Fig 4. a-b, Extended Data. Fig 5. a). Also, different T cell subclusters exert distinct effects around the ECM scaffold (Fig 4. d).

**Figure4.**
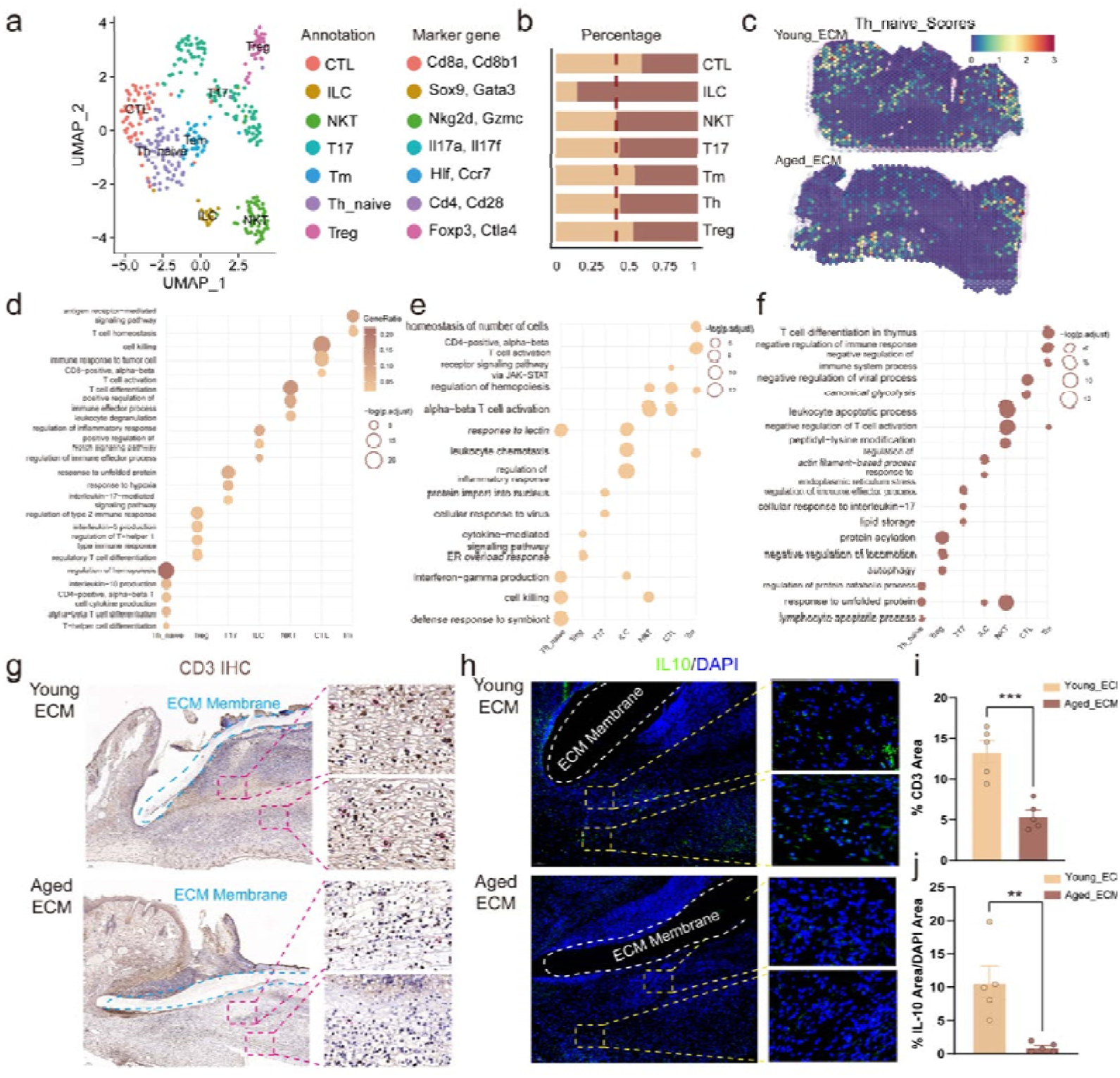
The intergroup difference of αβ T cells and the lack of type □ immune response in Aged_ECM. (a-b) The UMAP reduction results for the subclustering of αβ T cells. The annotations and marker genes are listed in the right. (c) The spatial distribution of crucial Th_naive subtype in type L immune response in spatial slices. (d) The representative GO enrichment results for each subtype of the wholeαβ T cells. (e-f) The upregulated GO enrichment terms of each subtype of T cells in Young_ECM and Aged_ECM, respectively. (g) The IHC staining for CD3 results shows more abundant adaptive immune cells in ECM mediated wound healing process in young mice. (h) The IF results for IL-10, the characteristic marker for type L immune response, shows its reduced expression around ECM scaffolds in aged mice. (i) The semiquantitative analysis for the CD3 positive area fraction and IL-10 immunofluorescence area around ECM scaffolds between groups (n=3 for each group). ***P<0.001 and **P<0.01by Student’s t-test for data in (i) and (j).

T17 secreting IL-17, Treg inhibiting immune hyperactivation, and ILC involved in tissue inflammation were identified and dispersed at distinct positions around the ECM scaffolds. Besides, Tm, primarily located in the fascia, upregulated genes such as hematopoietic stemness related Hlf^47^ and skin-homing related Sell^48^, and exhibited CD8+ T cell differentiation enrichment, implying their possible role as the CTL precursors from circulation. Moreover, NKT and CTL, as effector immune cells, although both demonstrated cytotoxic capabilities in clearing necrotic cells, but the spatial location are distinct with CTLs more abundant beneath the scaffold and papillary dermis in surrounding skin, while NKTs residing in the lower dermis in surrounding skin (Extended Data. Fig 5. d-i). Notably, Th_naïve exhibited upregulated differentiation potential to specialized T helper types, suggesting the precursor role as the T helper cells to coordinate the polarized immune responses through differentiation into different helper T cells (Th1, Th2, and Th17) and drive the type 1/2/3 paradigm of immunity^49^. The spatial location of Th_naïve were widely distributed, including the granulation tissue beneath the scaffold and surrounding skin (Fig 4. c), indicating their potential in shaping the immune microenvironment around the ECM scaffolds.

Afterwards, we explored the differential distribution and functions of these subtypes among aged and young groups (Fig 4. e-f, Extended Data. Fig 5. b). T17s displayed different intergroup spatial position to the ECM scaffold. T17s in Young_ECM are mostly distributed in the fibroblastrich granulation tissue below the ECM scaffold to enhance fibroblast migration and collagen deposition^50,51^, while located on the wound margin around scaffolds in Aged_ECM, where the secretion of IL-17 was proven to exacerbate local inflammation^52,53^, in line with the poor reepithelialization (Extended Data. Fig 5. f). In addition, Tm and CTL involved in cytotoxicity of adaptive immune response, showed significantly reduced abundance around the aged group scaffold, indicating compromised immune efficacy in clearing exogenous pathogens, which might contribute to the persistence of certain cell populations needing clearance, such as Mhsp, leading to prolonged local inflammation. Moreover, the Th_naïve downregulated Th2 related genes like IL-10 while upregulated genes linked to type I immune responses such as IL-1 in aged wounds, consistent with the differential intergroup spatial scores around the scaffold, suggesting the Th_naïve are more likely to differentiate in to Th1 rather than regeneration associated Th2 (Fig 4. g-j, Extended Data. Fig 5. c). Moreover, the downregulation ability to of endothelial regeneration and wound healing in Th_naïve has also been observed in Aged_ECM. the aged group scaffold downregulated its ability to regulate and wound healing, which further indicated the crucial role of Th_naïve in the disparities of the immune microenvironment around scaffolds between groups (Fig 5. e).

**Figure5.**
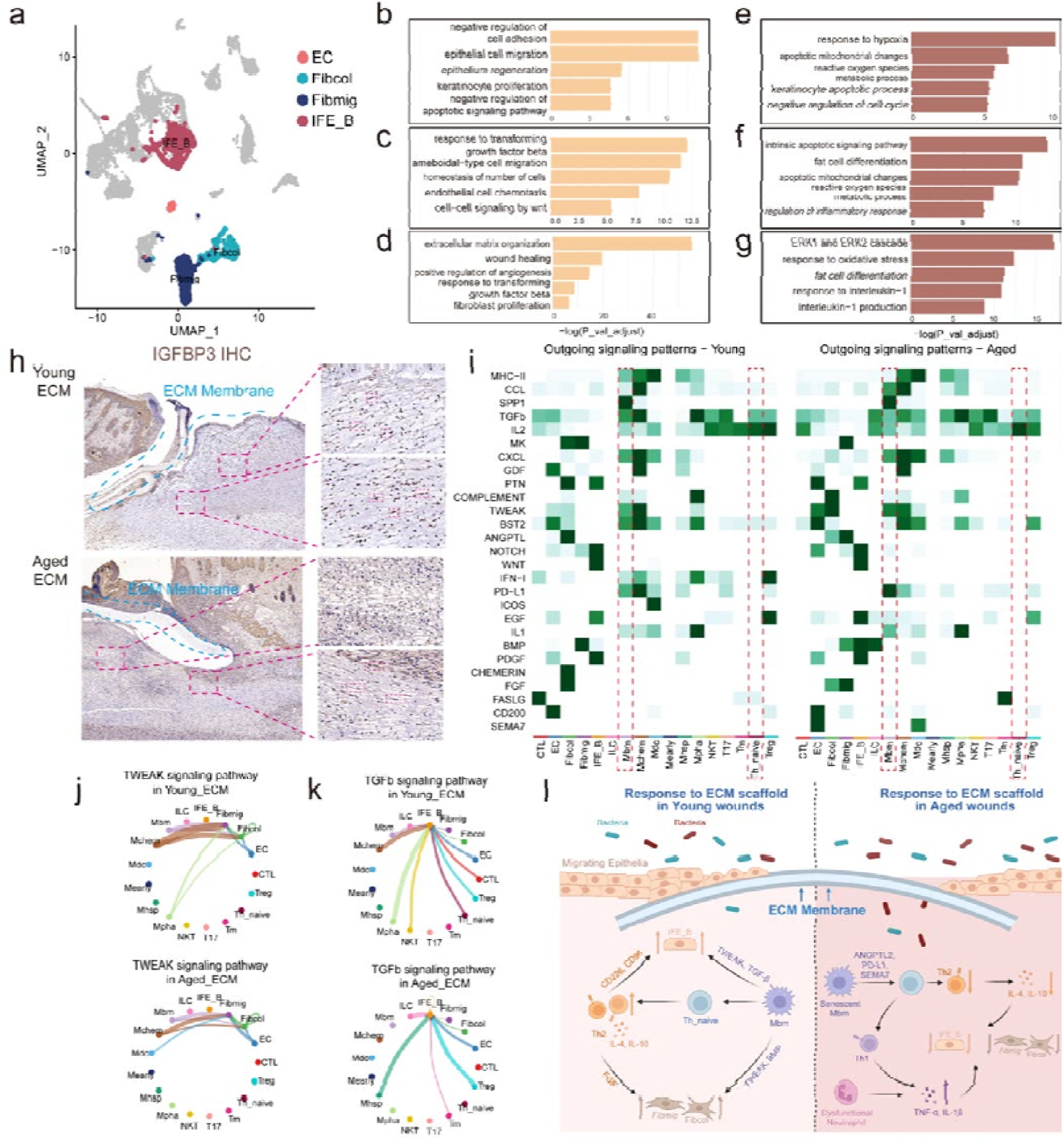
The cell-cell communication results account for the disparity of the performance in aged wounds. (a) The selected stromal cell types related to the delayed re-epithelization and dysregulated fibrous deposition in the first level UMAP reduction results. (b-d) The downregulated GO enrichment results in IFE_B, Fibmig and Fibcol respectively in Aged_ECM. (e-g) The upregulated GO enrichment results in IFE_B, Fibmig and Fibcol respectively in Aged_ECM. (h) The IHC staining results for IGFBP3 results revealed the slower migration ability of Fibmig in Aged_ECM. (i) The heatmap for the differential relative communication strength in total outgoing signaling pathway pattens between immune cells and stomal cells in Young_ECM and Aged_ECM. (j-k) The representative circle plots for the incoming pathway received by IFE_B and Fibcol respectively. (I) The schematic diagram for the difference of host defense and the effects of the microenvironment on the migrating IFE basal cells and fibroblasts around the ECM scaffolds.

Subsequently, through the gene set enrichment analysis of SenMayo, we identified CTL as the most notable subclusters affected by cellular senescence, consistent with previous reports^54^ while Th_naïve did not exhibit significant signs of cellular senescence^55^. This suggest the impairments in the cytotoxic capabilities of CTLs, may be, or at least in part, attributed to, the intrinsic cellular senescence, while the inability to differentiate into Th2 types and to build a regenerative microenvironment of Th_naïve is less likely the same reason but more likely due to different interactions with other cells around young and aged scaffolds.

### 4. Cell-cell communications elucidate the difference in the regenerative microenvironment and performance of ECM scaffolds between groups

Above all, we have identified the distinct functional subclusters of macrophages and T cells and the impact of their intergroup difference may have on the microenvironment. Besides, cellular interactions among immune cells may also contribute to the unique microenvironments responding to ECM scaffolds in different ages.

To begin with, we ran the secondary dimension reduction based on the first level reduction result and focused the subclusters related to epidermal and dermal repair respectively, namely IFE_B (Krt5+, Col17a1+), Fib_mig (Igfbp3+, Cebpd+) as well as Fib_col (Col1a1+, Col1a2+) (Fig 5. a, Extended Data. Fig 6. a). IFE_B was considered as the epidermal stem cell pools to migrate and proliferate in response to wounds. Although previous results demonstrated the stemness of IFE_B were not affected during aging^56,57^, IFE_B exhibits challenges in the migration and proliferation, in accordance with the slower re-epithelialization in Aged_ECM (Fig 5.b, e). Similarly, Fib_mig and Fib_col with the capacity for extracellular matrix synthesis and collagen deposition, showed deceased migration (Fig 5.h) but more tendency to the adipocyte, in line with the dermal repair in Aged_ECM (Fig 5. c-d, f-g). We then merged these cells with previously identified subclusters of macrophages and αβ T cells to speculate the interaction patterns and intergroup differences.

Cell-cell communication revealed alterations of incoming and outgoing pathways were most significant in Mbm, further demonstrating its susceptibility to aging around the scaffold (Extended Data. Fig 6. f). As for the outgoing pathways of Mbm (Fig 5.i, Extended Data. Fig 6. h-j), a more enriched pro-inflammatory IL-1 pathway was discovered in aged group, which could potentially lead to chemotaxis for more Mpha and acute inflammatory cells, in line with the observations. Besides, the interaction between Mbm and Th_naïve in Aged_ECM also helps explain the impaired type L immnue response around the scaffold. On one hand, decreased CXCL chemotaxis pathways by Mbm correlated with lower abundance of Th_naïve cells around scaffold^58^. On the other, downregulation of MHC-II molecules and upregulation of immunosuppressive PD-L1 pathways in Mbm accounted for the reduced co-stimulatory signals and activation of Th_naïve cells^59^. In accordance, the signal strength of Th2 related IL-10 pathway was weaken even in fewer interactive cell types. As for the incoming pathways of Mbm (Extended Data. Fig 6. g), ANGPTL2^60^ and SEMA7^61,62^ pathways from a wide range type of cells in aged group have been proven to facilitate the pro-inflammatory, rather than pro-healing macrophage phenotype shift. Additionally, the more enriched CD200 pathway^63^ in aged Mbm may also explain the less production of subsequent IL-10. Besides, local Fib_mig and EC in the Aged_ECM group not only lost the ability of macrophage chemotaxis via the declining CHEMERIN pathway^64^, but also failed to promote the antiinflammatory macrophage phenotype shifts through downregulation of MK and BST2 pathway^65,66^. Therefore, the multiple incoming and outgoing pathways of M2bm collectively contribute to the impediment in local regenerative microenvironments.

After exploring the difference in the communication pathways between groups, we also analyzed the interaction between immune cells and stromal cells to for better understanding the poor performance of ECM scaffolds in wound healing. In aged group, although Mchem and Fib_col expressed active PTN^67^ and FGF^68^ pathways related to epithelial migration and proliferation towards IFE_B, Mbm around the scaffold showed lower strength of WNT^69,70^ and GDF^71^ pathways, which seems to dominate the IFE_B proliferation and migration activities, associated with delayed re-epithelialization. Conversely, Mbm in Young_ECM exhibited more activity through the TWEAK pathway with IFE_B (Fig 5. j), previously known to promote more chemokine secretion by epithelial cells, contributing to the establishment of regenerative microenvironment in turn^72^. As for dermal repair, BMP, PDGF, FGF, and TWEAK ^73–76^ pathways which are shown to promote the proliferation and activation of fibroblast (Fig 5. k), were downregulated by Mbm and EC cells, in line with the poor and irregular collagen deposition and tissue repair after implantation.

Therefore, the immune cells and non-immune cells communication helps explain the reason for the persisting acute inflammation and poor performance of ECM scaffold in aged wounds through differential pathway activities (Fig 5. l).

## Discussion

Advances in technology lead to the design and application of numerous biomaterials to better wound healing issues, based on various concepts and mechanisms. However, most attention has been focused on the performance in view of the structure or components of these materials, the subsequent reactions post-implantation into diverse organisms are scarcely addressed^77^. In the context of different individuals with altered pathophysiological environments, the same material might exhibit distinct characteristics, which might be the reason for the failure of the clinical practice in some patients^78^. In this study, we observed the remarkable facilitation of early wound closure and later HF neogenesis by ECM scaffolds in young wounds. ECM scaffolds also achieved the dermal repair results with more regular and denser fiber repair and vascular regeneration. Curiously, the same ECM scaffolds into aged wounds failed to promote wound healing process in epidermis as well as dermis. Therefore, based on the principles in precision medicine, exploring the reasons for differential performance of ECM scaffolds in aging individuals is necessary for future adjustments.

However, the most challenge encountered when thoroughly exploring mechanisms of biomaterials post-implantation is the coexistence of multiple cell types with distinct functional statuses and different locations^24,46^. To solve this, we integrated scRNA-Seq with spatial transcriptomics, and successfully identified various functional statuses at single-cell resolution as well as their respective spatial niches around the ECM scaffolds, elucidating the detailed biomaterial-host interaction mechanism. Unlike the M1/M2 classification system, the integrated analysis reveals six states based on the functional similarity, which scatter and respond to the ECM scaffolds in spatial splices as mentioned above. Among them, particular attention was directed towards the *Spp1+Arg1+*Mbm, beneath and displaying the strongest interaction with the ECM scaffold, with various functions such as antigen presentation, cell chemotaxis, and regulation of adaptive immune response. Furthermore, we identified the undiscovered role of Mbm to activate Th_naïve cells and Th2 differentiation, fostering a regenerative microenvironment to enhance the functional wound healing, which is missing around the ECM scaffolds in aged wounds.

We then tried to explain the mechanisms of the altered function and response to ECM scaffolds between groups in different statuses of macrophage and T cells. As with the intrinsic aging process, cellular senescence is responsible for the functional declination in many cells, but the dentification of cellular aging at the transcriptomic level may not be enough based on the current enrichment databases^42^. Therefore, we turned to the Senmayo gene set^41^ which exhibited the excellent evaluation for cellular senescence and observed significant senescent signatures exclusively in Mbm cells, which helps explain the functional defects around scaffolds in aged wounds. However, type L immune response-associated CD4+ Th_naïve did not show the cellular senescence^79^, and its functional defects may be attributed to the local cell-cell communications with corresponding Mbm. Through CellChat analysis, the differential pathway expression from Mbm to Th_naïve further supported the initial speculation. Finally, we also unveiled the crosstalk between immune cells and stromal cell related to wound healing and provided a comprehensive understanding of the disparity of host defense to and performance of ECM scaffolds in aged individuals.

## Methods

### 1. Ethical approval

All procedures were approved by the Institution Review Board of West China Hospital of Stomatology (WCHSIRB-D-2023-294).

### 2. Fabrication of ECM scaffolds

The ECM scaffold membrane of aligned fibers with approximate 300 nm diameter were fabricated using the electrospinning technology. 20% w/v poly (lactic-co-glycolic-acid) (PLGA) (LA/GAL=L75:25, molecular weight = 105 kDa, Jinan Daigang Biomaterial Co. Ltd.) and 2% w/v fish collagen (FC) (Sangon Biotech Co. Ltd.) were completely dissolved in 1,1,1,3,3,3-Hexafluoro2-propanol (HFIP) (Aladdin Co., Ltd.) with stirring. The solutions were then loaded into a sterile syringe with a flat-tipped 21 G stainless steel needle. When electrospinning, the voltages of -2/5kv and 11 cm distance between the needle and the rotating cylinder (2800Lrpm) were used. After collected and dried in a vacuum oven for 48h, the morphology of PLGA/FC ECM scaffold was observed by scanning electron microscopy (SEM; JSM-7500F, JEOL, Japan). Afterwards, the scaffolds were cut into circular shapes (8mm) and sterilized with γ-irradiation before the animal implantation experiments.

### 3. Full-thickness wound model and implantation procedures

All procedures involving animals were approved by the Institution Review Board of West China Hospital of Stomatology (WCHSIRB-D-2023-294). Young (7-week-old) and aged (88-weekold) male C57BL/6J mice (Dossy Experimental Animals Co., Ltd.) were used in this research. The mice were housed under standard conditions including temperature of 21–27 °C, humidity of 40– 70%, and a 12 h light-dark cycle with free access to food. The number of animals used for each experiment is indicated in the figure legends. To mimic the wound healing process in human, the mice-splinted model was conducted to minimize wound contraction by the panniculus carnosus of rodents^80^. The circular (diameter = 6mm) full-thickness wounds were created in the mice dorsal skin and stented by silicone loops. The mice in further study were divided into four groups: young mice treated with normal saline (Young_Ctrl) or ECM scaffolds (Young_ECM) below the wound, and aged mice treated with normal saline (Young_Ctrl) or ECM scaffolds (Young_ECM) below the wound. Afterwards, the dorsal wounds were covered with sterile Tegaderm film (3M) and fixed on the silicone ring. Mice were euthanatized at 1–4 weeks after the implantation, and the full-thickness sample was harvested.

### 4. Single cell RNA sequencing

#### 4.1 Sample Preparation and Single-Cell Isolation

Three fresh samples were collected POD7 respectively in Young_ECM and Aged_ECM group. Briefly, the wound samples were enzymatically digested by the Epidermis Dissociation Kit (Epidermis Dissociation Kit, mouse; Miltenyi Biotec) to achieve the epidermal-dermal separation. After that, the epidermis was then dissociated by a gentleMACS Dissociator (Miltenyi), filtered (70mm cell strainer, Corning, Durham), centrifuged (300 g, 10 min, 4 °C), and resuspended with phosphate-buffered saline (PBS) containing 0.5% bovine serum albumin (BSA) while the dermis was cut into 0.5 mm width pieces, treated with pre-mixed enzyme solution containing type I collagenase (Gibco, Grand Island) and trypsin (Gibco, Canada), dissociated by gentleMACS Dissociator (Miltenyi), and digested for 2.5 hours in a hybridization oven (Peqlab PerfectBlot). After being dissociated, filtered, centrifuged, and resuspended in red blood cell lysis buffer (Solarbio), the dermis cells were mixed with the epidermis cells. Then Dead Cell Removal MicroBeads (Miltenyi) was used to remove the dead cells and debris.

#### 4.2 Sequencing and data processing

Single-cell suspensions were then carried out for Single-Cell RNA-seq (10x Genomics Chromium Single Cell Kit). Sequencing (10x Genomics Chromium Single Cell 3′ v3) was performed using an Illumina 1.9 mode. Then, reads were aligned, and expression matrices were generated for downstream analysis (Cell Ranger pipeline software).

#### 4.3 Downstream computational analysis

First, ambient RNA signal was removed using the default SoupX (v1.4.5) workflow^81^. Young and aged samples were then merged into one Seurat object and preprocessed using the standard Seurat (v4.1.0) workflow (NormalizeData, ScaleData, FindVariableFeatures, RunPCA, FindNeighbors, FindClusters, and RunUMAP). After preprocessing, DoubletFinder (v2.0) was used to identify putative doublets in each dataset, individually. Estimated doublet rates were computed by fitting the total number of cells after quality filtering to a linear regression of the expected doublet rates published in the 10x Chromium handbook. Then, the integration of each sample and future analysis was carried out in Seurat (v4.1.0). RunPCA function was used to process the most variable genes determined by the FindVariableGenes function (selection.method = \“vst\“, nfeatures = 2000). ElbowPlot function was performed to determine the number of principle components input. RunUMAP function (Seurat) with the first 18 principal components as input was performed for dimensionality reduction. Unsupervised clustering was performed using the FindClusters function of Seurat and clustree R package (v0.5.0) and differentially expressed genes were determined by the FindAllMarkers and FindMarkers function. ClusterProfiler (v4.6.2) was used to perform the gene set enrichment analysis and the visualization was performed by GseaVis (v0.0.5). CellChat (v1.5.0) was used to predict receptor-ligand probability among cell subpopulations.

### 5. Spatial transcriptomics

#### 5.1 Slide preparation

The wound samples in Young_ECM and Aged_ECM group were collected immediately after harvesting and fixed overnight in 4% neutral buffered formalin for the FFPE protocol. Tissues underwent three 5 min washes in PBS at room temperature followed by dehydration washes in increasing ethanol concentrations. After dehydration, tissue was processed using a Leica ASP300 Tissue Processing for 1 hr. Tissues were then embedded in paraffin and stored at 4°C. Then FFPE samples were cut at the standardized 5 µm thickness onto the 6.5 mm × 6.5 mm capture areas. Slides then underwent deparaffinization and H&E staining according to the manufacturer’s instructions (10X Genomics, Visium Spatial). Decrosslinking was performed to release RNA that was sequestered by the formalin fixing, followed by probe hybridization, ligation, release, and extension according to the manufacturer’s instructions. Finally, FASTQ files and histology images were processed (SpaceRanger software) for genome alignment.

#### 5.2 Downstream dimension reduction analysis

Raw output files for the sample of Young_ECM and Aged_ECM were read into R studio with the Seurat R package (v4.1.0). Normalization across spots was performed with the SCTransform function. The clustering results were annotated based on the dimension reduction results and anatomic structure.

#### 5.3 Integration analysis of scRNA-seq and ST

Signature scoring derived from scRNA-seq and ST signatures was performed with the AddModuleScore function in Seurat R packages (v4.1.0)^82^. Firstly, the FindAllMarkers function was performed to identify the top 20 marker genes (based on avg_log2FC value) of each cluster in scRNA-seq results. AddModuleScore was then used to calculate the average scoring of each gene set in every spot of the spatial transcriptome. Finally, the scores were mapped to the spatial transcriptome using the SpatialFeaturePlot function.

### 6. R analysis packages

R v4.2.3 was used for downstream analysis of single-cell RNA sequencing and spatial transcriptomics data. R packages used: Seurat(v4.1.0), harmony (v0.1.1), clustree (v0.5.0), tidyverse (v2.0.0), Matrix (v2.0.0), ggplot2 (v3.4.2), DEseq2 (v1.32.0), GOplot (v1.0.2), stringr (v1.5.0), EnhancedVolcano (v1.10.0), ClusterProfiler (v4.6.2), org.Mm.eg.db (v3.13.0), Cellchat (v1.5.0), patchwork (v1.1.2), data.table (v1.14.8), hdf5r (v1.3.8), pracma (v2.4.2), GseaVis (0.0.5). The tutorial of Seurat software is available at https://satijalab.org/seurat. The tutorial of ClusterProfiler software is available at https://github.com/YuLab-SMU/clusterProfiler. The tutorial on Cellchat software is available at https://github.com/sqjin/CellChat.

### 7. Histopathology, Immunohistochemistry, and immunofluorescence microscopy

The samples were fixed with 4% paraformaldehyde at least 48 hours before ethanol and xylene dehydration. H&E staining and Masson’s trichrome staining were performed for the observation of re-epithelialization and collagen fiber deposition. Immunohistochemistry staining for CD31 (280831-AP, Proteintech, 1:1000) and Igfbp3 (10189-2-AP, Proteintech, 1:100) were performed for to assess the angiogenesis and fibroblast migration respectively. For the evaluation the acute inflammatory molecules and immune cells infiltration, immunohistochemistry staining for CD3 (14-0032-82, Thermo Fisher Scientific, 1:100) and immunofluorescent staining for Ly6g (65140-1Ig, Proteintech,1:200), iNOS (22226-1-AP, Proteintech,1:200), IL-1β (P420B, Thermo Fisher Scientific, 1:200),TNF-α (60291-1-Ig Proteintech, 1:200), OPN (22952-1-AP, Proteintech, 1:200), F4/80 (29414-1-AP, Proteintech, 1:200) and IL-10 (ARC9102, Thermo Fisher Scientific, 1:100) were performed.

## Acknowledgments

We appreciate the Novogene company for scRNA-seq and ST sequencing work, and the National Clinical Research Center for Oral Diseases & State Key Laboratory of Oral Diseases for animal and histological experiments, the Analytical & Testing Center of Sichuan University for the fabrication of ECM scaffolds, and the BioRender website for the illustration of the schematic diagrams. We also thank Chen Hu, Xin-hui Li, Zu-mu Yi, and Tian-tian An for the help to this work.

## Funding

National Natural Science Foundation of China grant 81970965; 82271015 (Y.M.)

Interdisciplinary Innovation Project, West China Hospital of Stomatology Sichuan University grant RD-03-202006 (Y.M.)

Research and Develop Program, West China Hospital of Stomatology Sichuan University grant LCYJ2019-19 (Y.M.)

## Author contributions

Conceptualization: SDC, CCY, YM

Methodology: SDC, CBW

Investigation: SDC

Visualization: SDC

Writing – original draft: SDC

Writing – review & editing: CCY, YM

Project administration: SDC

Funding acquisition & Supervision: YM

## Competing interests

Authors declare that they have no competing interests.

## Data and materials availability

All data to support the conclusions in this work can be found in the figures and the Extended Data. other data can be requested from the corresponding authors.

---

**Extended Data. Fig 1.**
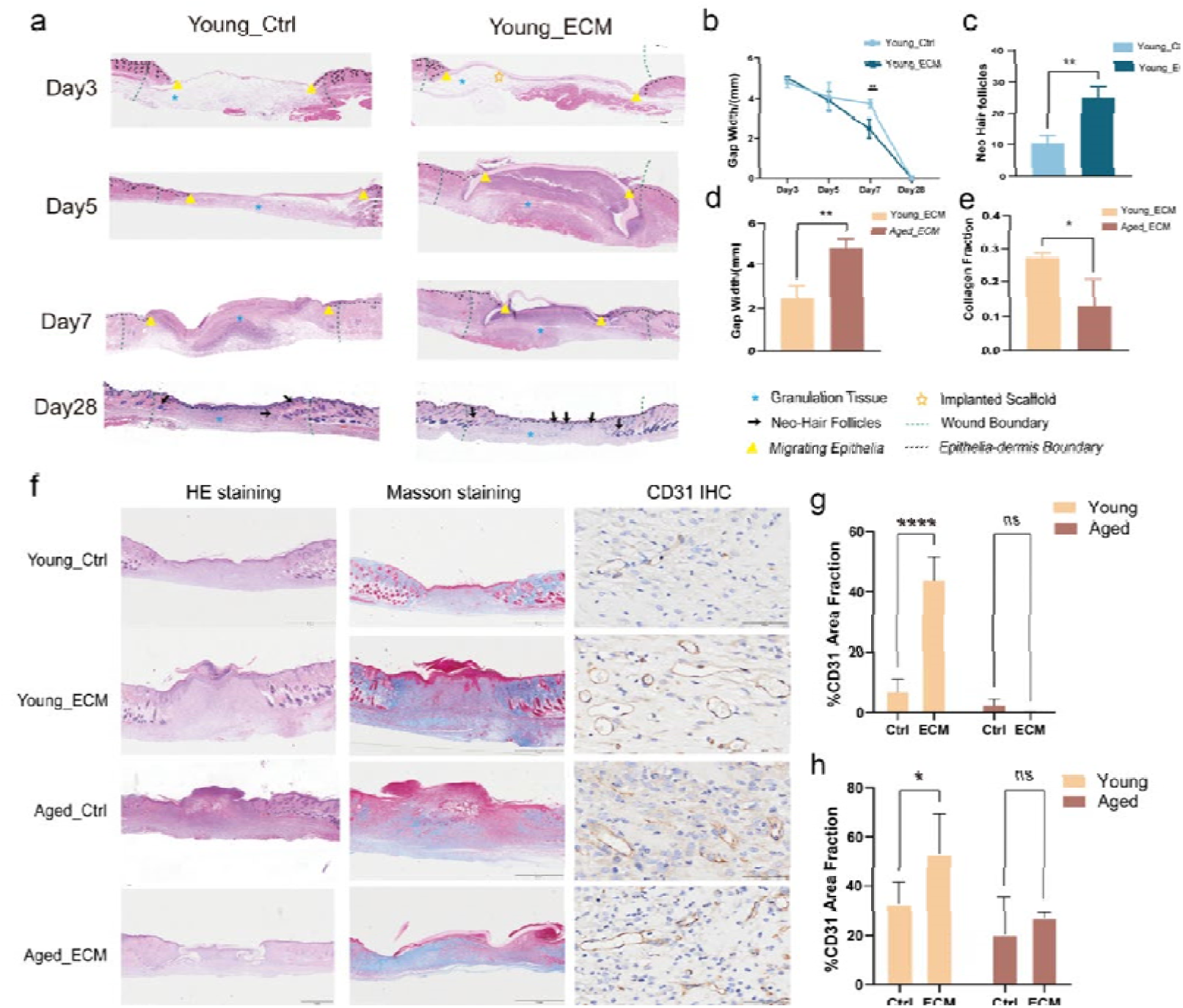
The outperformances of ECM scaffolds in young mice. (a) The histologic H&E results for the ECM group and Control group in young mice at POD3, 5, 7 and 28 days. (b) The semiquantitative results of gap width of Control group and ECM group in young mice. (c) The quantitative results of hair follicle neogenesis of Control group and ECM group in young mice at POD28. (d-e) The comparison of the gap with and collagen deposition in ECM groups in young and aged mice. (f) The histological results for H&E as well as Masson staining and the IHC results for CD31 of wounds at POD14 in Control group and ECM group of young and aged mice. (g-h) The semiquantitative analysis of CD31 area fraction in different groups at POD7 and POD14 respectively.

**Extended Data. Fig 2.**
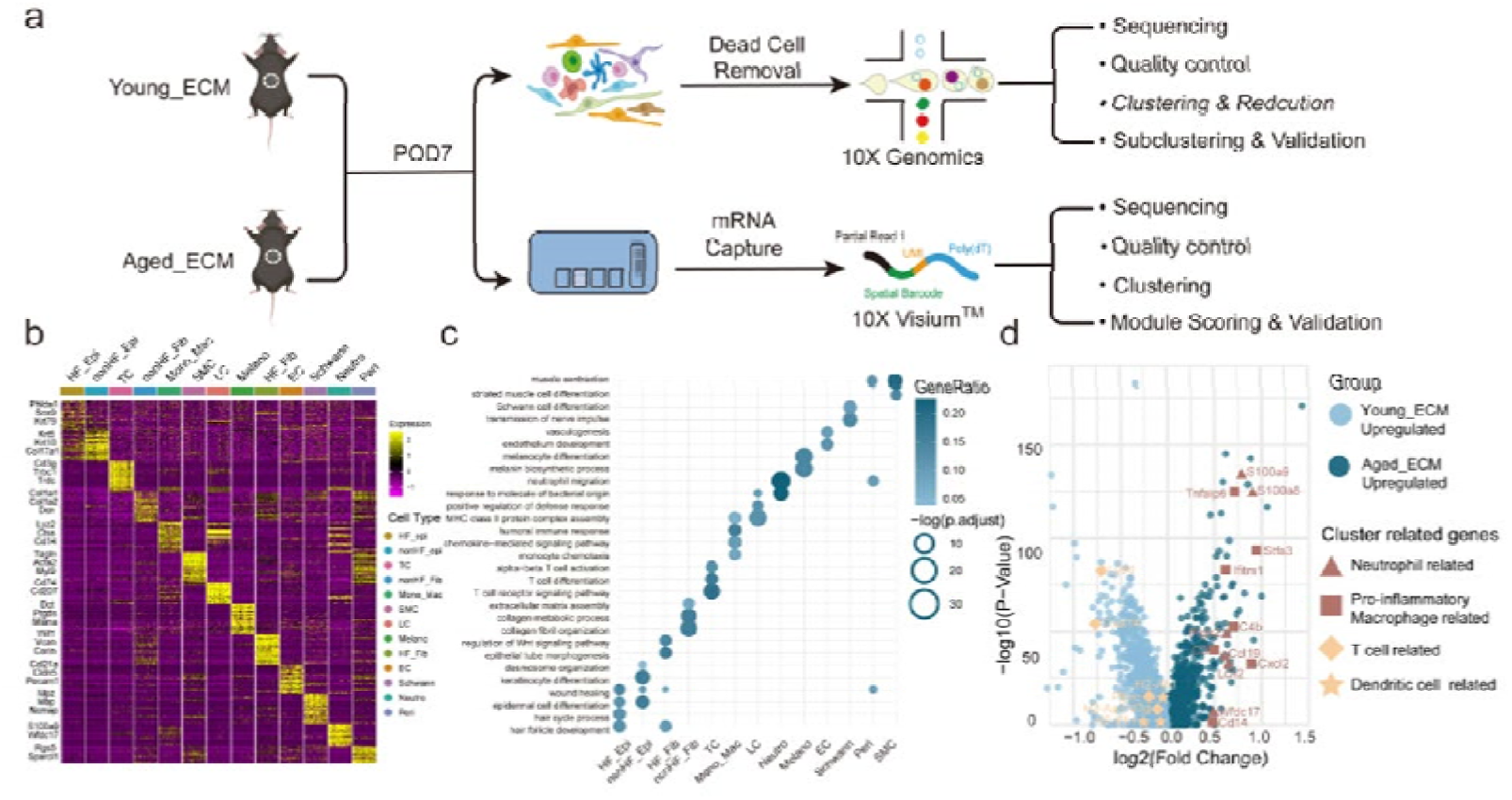
The first-level reduction results and differential expressed genes (DEGs) between groups. (a) The workflow for scRNA-seq and spatial transcriptomics. (b) The heatmap for top20 marker genes of respective cell types in the first level reduction. (c) The representative GO enrichment results for each cell type in the first level reduction. (d) Volcano plot showing interested differentially expressed genes in Young_ECM and Aged_ECM.

**Extended Data. Fig 3.**
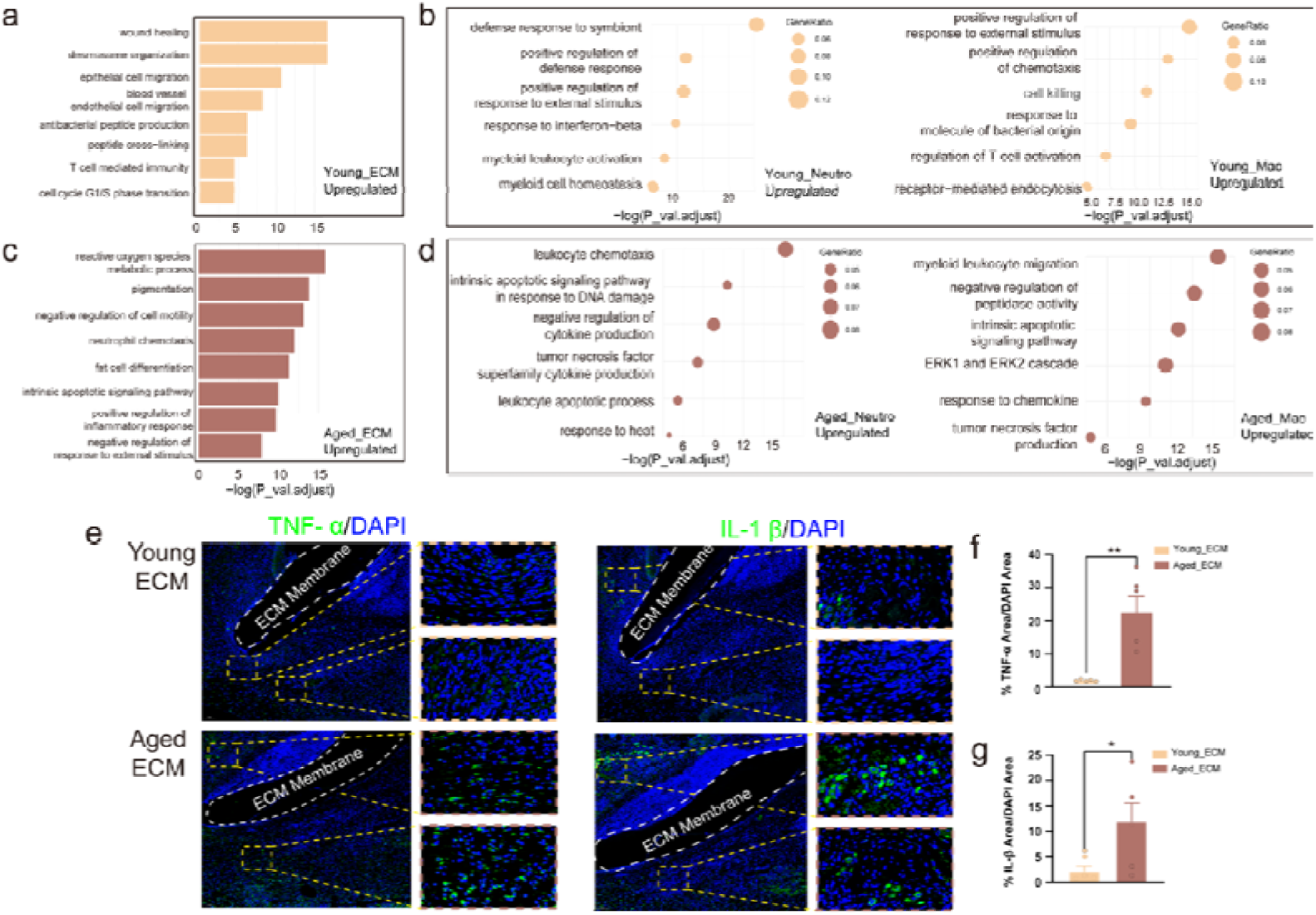
The functional intergroup difference in pseudobulk results and acute inflammation related cell types. (a,c) The downregulated GO enrichment results and upregulated results of Aged_ECM in all types of cells in the wounds. (b) The downregulated GO enrichment results for neutrophils (left) and macrophages (right) in Aged_ECM. (d) The upregulated GO enrichment results for neutrophils (left) and macrophages (right) in Aged_ECM. (e) The IF results for TNF-α as well as IL-1β, the characteristic marker for acute immune response, shows its higher expression around ECM scaffolds in aged mice. **P<0.01 and *P<0.05by Student’s t-test for data in (e).

**Extended Data. Fig 4.**
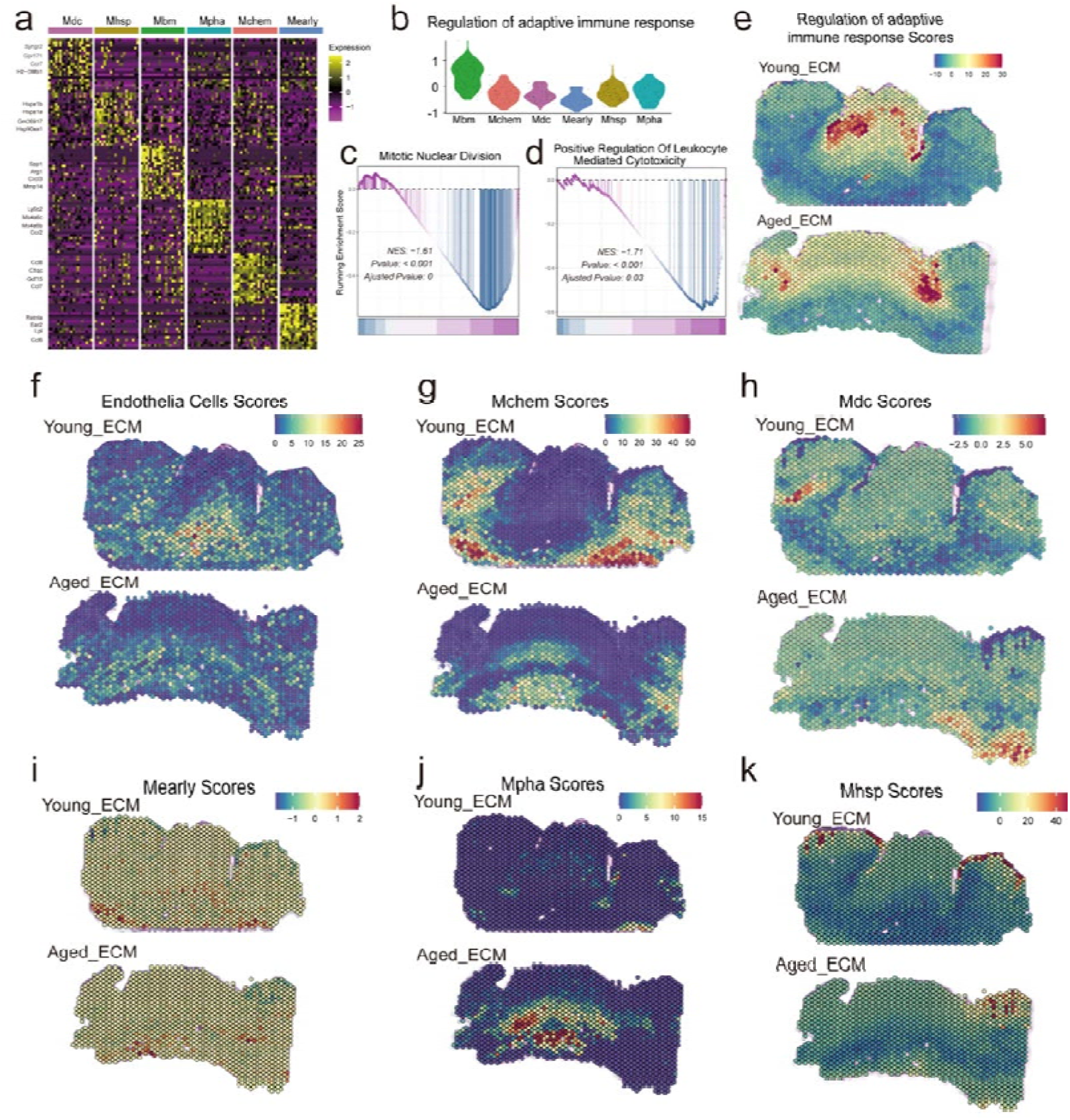
The spatial distribution of subtypes of macrophages and their intergroup difference. (a) The heatmap plot showing the top 20 DEGs for each subtype of the macrophages. (b) The violin plot showing the AddModule scores results for the term “regulation of adaptive immune response” in each subtype of macrophages. (c-d) The downregulated GSEA enrichment results for Meraly and Mdc in Aged_ECM. (e) The spatial scores for the term “regulation of adaptive immune response” in Young_ECM and Aged_ECM. (f-k) The AddModule scores for the top20 DEGs for Endothelial cells, Mchem, Mdc, Meraly, Mpha and Mhsp to show their spatial distributions respectively.

**Extended Data. Fig 5.**
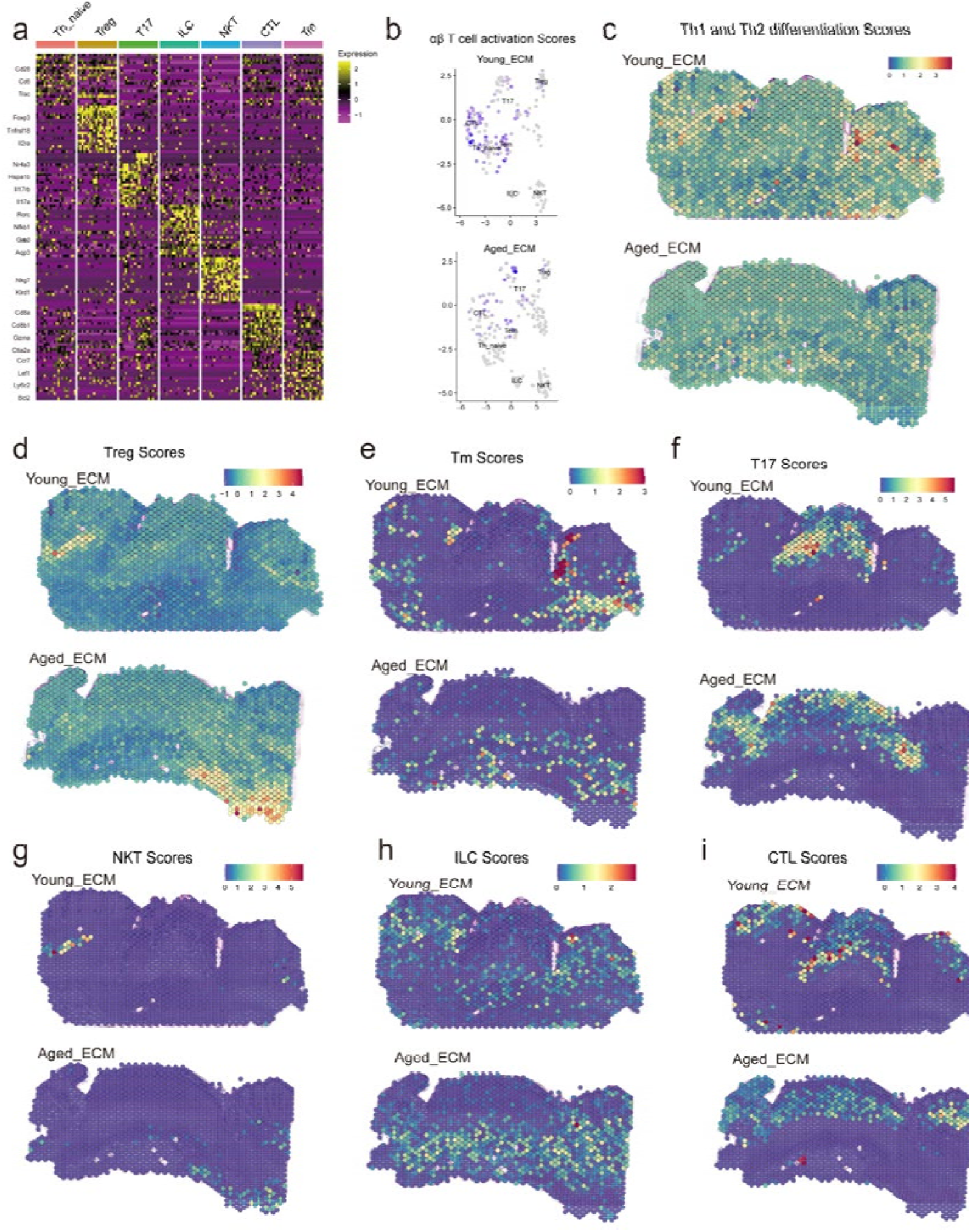
The visualization for the subclustering of αβ T cells and their spatial distributions. (a) The heatmap plot showing the top 20 DEGs for each subtype of the αβ T cells. (b) The FeaturePlot t showing the AddModule scores results for the term “αβ T cell activation” in the UMAP subclustering reduction result. (c-d) The spatial scoring results for the downregulated KEGG enrichment result of scRNA-seq in Aged_ECM. (d-i) The AddModule scores for the top20 DEGs for Treg, Tm, T17, NKT, ILC and CTL to show their spatial distributions respectively.

**Extended Data. Fig 6.**
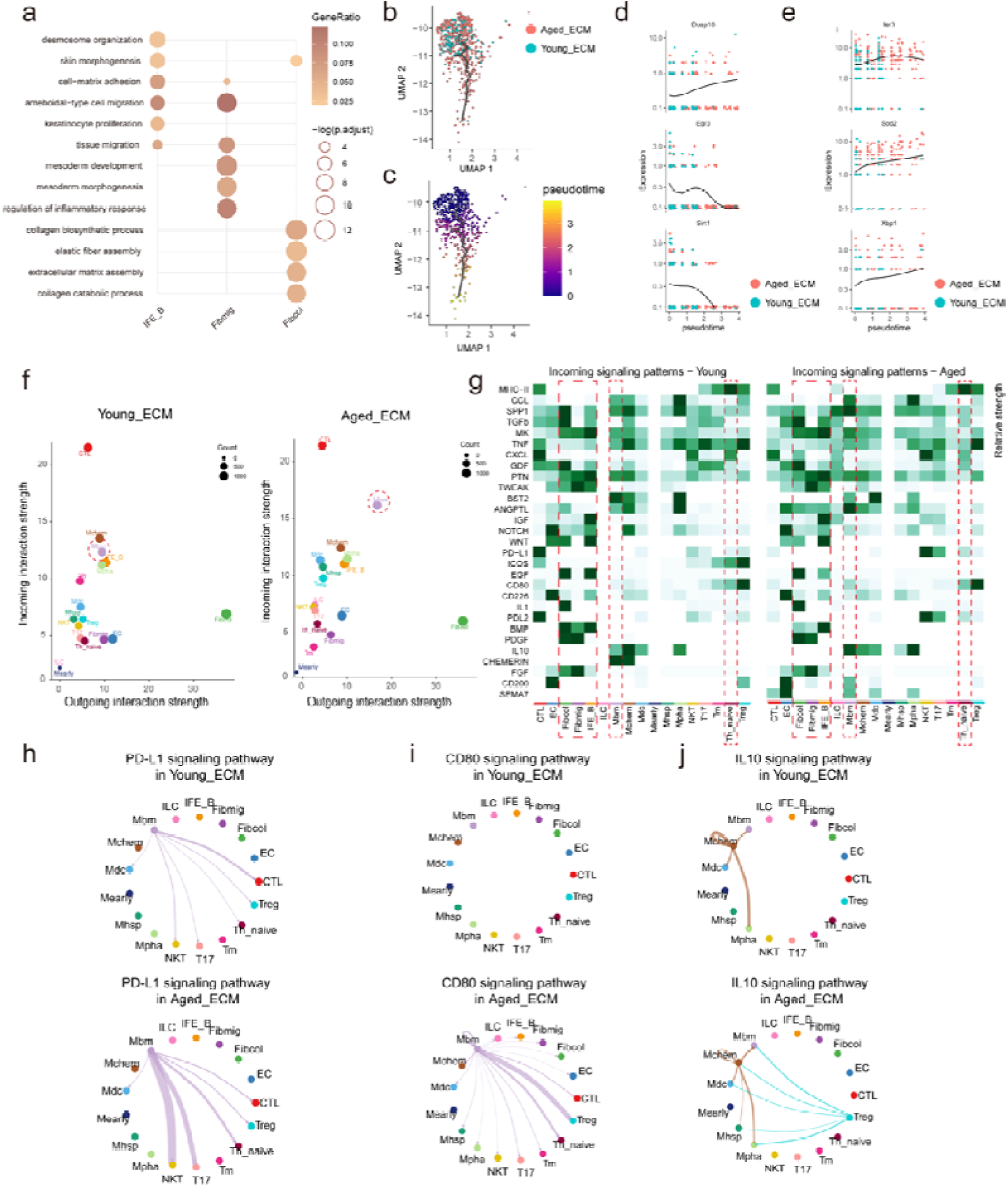
The gene enrichment results for stromal cells and the cell communications results in immune cells and stromal cells. (a) The GO enrichment results for each stromal cell type in Fig5a. (b-c) The selected UMAP reduction results and pseudo-time trajectory in Monocle3 for Fibmig. (d-e) The relative gene expression along pseudo-time trajectory to show their intergroup difference associated with the enrichment terms “tissue migration” and “intrinsic apoptosis pathways” respectively. (f) The role of different types in the outgoing and incoming pathways in Young_ECM and Aged_ECM. (g) The heatmap for the differential relative communication strength in total incoming signaling pathway pattens between immune cells and stomal cells in Young_ECM and Aged_ECM. (h-j) The representative outgoing pathways secreted by Mbm and Th_naive in Young_ECM and Aged_ECM in accordance with the disparities around the ECM scaffolds.

## Notes

### Competing Interest Statement

The authors have declared no competing interest.

